# *In silico* description of a new Amalgavirus in the *Cannabis sativa* transcriptome

**DOI:** 10.1101/2025.02.27.640618

**Authors:** Matheus de Castro Leitão, Fernando Lucas Melo

## Abstract

*Cannabis sativa* has a variety of industrial interest products, such as phytocannabinoids, terpenoids, phenylpropanoids, acyl sugars, and fibers. Several described pathogens, including viral species, will impact the current green cannabis revolution. The recent sequencing of its genome and transcriptome, allowing the optimization and understanding of the production of the metabolite, are a relevant tool for viruses presence analyzing. Using the cannabis transcriptome and Data mining analysis, we describe the first amalgavirus infecting Cannabis, the Cannabis sativa amalgavirus 1 (CSA1). The plant amalgaviruses has nine species infecting relevant crops. Like the other genus members, this cannabis virus has approximately 3.5 kb with two partially overlapping putative open reading frames with the characteristic +1 programmed ribosomal frameshifting. Mainly detected in the male plant, the CSA1 mapped reads were present in the flower and leaf tissues. Nevertheless, the possible impacts of viral replication on host metabolism and the production of secondary compounds are unknown.

---

Despite its domestication and use of at least 4,000 years B.P. (Long et al., 2017; McPartland et al., 2018; Ren et al., 2021), Hemp (Cannabis sativa L.) has only recently gained notoriety for its unique characteristics. Among them, there is the production of exclusive secondary metabolites, the phytocannabinoids, such as cannabidiol (CBD) and delta-9-tetrahydrocannabinol (THC) (Gülck and Møller, 2020; Hanuš et al., 2016), which have several pharmacological applications (Bonini et al., 2018; Page et al., 2020). Glandular trichomes mainly synthesize these compounds in high density in the floral buds of the female plant (Kim and Mahlbeg, 1997; Stout et al., 2012; Gülck and Møller, 2020). Other relevant products, with different properties, are terpenoids, phenylpropanoids, acyl sugars, and fiber (Andre et al., 2016). With an increase in its cultivation in several countries, whether for recreational or medicinal use, several projects have been sequencing its genome (van Bakel et al., 2011; Gao et al., 2020; Grassa et al., 2021), transcriptome (Braich et al., 2019), and reconstructing its domestication history (Ren et al., 2021) to optimize and understand the production of these metabolites.

As with other crops, the budding green revolution of hemp will probably be impacted by phytosanitary issues that can affect its production and alter essential metabolic characteristics (Fike, 2016; Welling et al., 2016). Several pathogens have already been described, including fungal, bacterial, nematode, and viral species (Chiginsky et al., 2022; Punja, 2021). The description of Cannabis virus infections is aged, dating back to the 1940s (Röder, 1941). Under experimental conditions, hemp is susceptible to agents known to infect other relevant crops, such as potato virus Y (PVY), cucumber mosaic virus (CMV), alfalfa mosaic virus (AMV), and others (Kegler and Spaar, 1997). In plantations, the reports are more recent and include hop latent viroid (HLVd) (Bektas et al., 2019; Warren et al., 2019), lettuce chlorosis virus (LCV) (Hadad et al., 2019), beet curly top virus (BCTV) (Giladi et al., 2020; Hu et al., 2021;), and new isolates cannabis cryptic virus (CanCV) (Ziegler et al., 2012; Righetti et al., 2018) and cannabis sativa mitovirus 1 (CasaMV1) (Nibert et al., 2018).

Recently identified, the *Amalgaviridae* family has two consolidated genera, *Zybavirus*, with a species that infects yeast (Depierreux et al., 2016) and *Amalgavirus* (Adams et al., 2014; Liu and Chen, 2009; Martin et al., 2011; Sabanadzovic et al., 2009, 2010), with 9 species that infect plants, and a proposed genus, “Anlovirus” associated with entomopathogenic fungi, giant springtails, and double-ended bristles (Pyle et al. 2017). Members of the amalgavirus genus, known as plant amalgaviruses, have single-segment, positive-sense, double-stranded RNA (dsRNA) genomes ranging from 3,427 to 3,437 bp (Liu and Chen, 2009; Martin et al., 2011; Sabanadzovic et al., 2009, 2010). The genome has two partially overlapping putative open reading frames (ORFs), believed to encode two proteins. The ORF1 product has an unknown function and may be a filamentous nucleocapsid protein (Krupovic et al., 2015) or a replication factory matrix-like protein (Isogai et al., 2011). The second protein is a fused product encoded by ORF 1 + 2 that is translated consequential to +1 programmed ribosomal frameshifting (PRF) where the portion of ORF2 has conserved motifs from viral RNA-dependent RNA polymerase (RdRp) (Liu and Chen, 2009; Martin et al., 2011; Sabanadzovic et al., 2009, 2010). Its genomic organization resembles members of the Totiviridae and because they have phylogenetic relationships with the Partitiviridae, it is proposed that amalgaviruses represent a link between these two families (Krupovic et al., 2015; Sabanadzovic et al., 2009). Amalgaviruses are vertically transmitted through seeds, and the formation of *bona fide* virions has not yet been detected, nor associated with symptoms in individual infections (Liu and Chen, 2009; Martin et al., 2011; Sabanadzovic et al., 2009, 2010).

In this work, we described a new Amalgavirus genomic sequence on the published transcriptome assembly generated by RNA-Seq (Braich et al., 2019) (Figure 1). The transcripts were first filtered using the Kaiju online version for taxonomic classification (Menzel et al., 2016). Using the Genome Detective Virus Tool (Version 1.130) (Vilsker et al., 2019), these subset dsRNAs were screened to merge the same ones, exclude host hits sequences, and look for similarities. All sequences with hits matching the viral database were then subjected to the NCBI BLASTx search against the complete nr database to exclude false positives. To confirm the assembly results and further extend incomplete genomes, trimmed reads were mapped back to the viral contigs and reassembled until genome completion or no further extension. The final contigs were annotated using the Geneious program (v. 9.1.3, Biomatters, Auckland, New Zealand). We selected a solid similarity contig to Phalaenopsis equestris amalgavirus 1 to further analysis. The open read frames were annotated using the following parameters: 500 bp minimum size, standard genetic code, and ATG start codon. Additionally, we used the UUU_CGN as a +1 ribosomal frameshifting motif prevalent among plant amalgaviruses to determine the two genomic ORFs. All the virus genome sequences had significant coverage, primarily in the male plant (flower and leaf), with 70,000 reads mapped in both individuals (Figure 1).

**Fig. 1.**
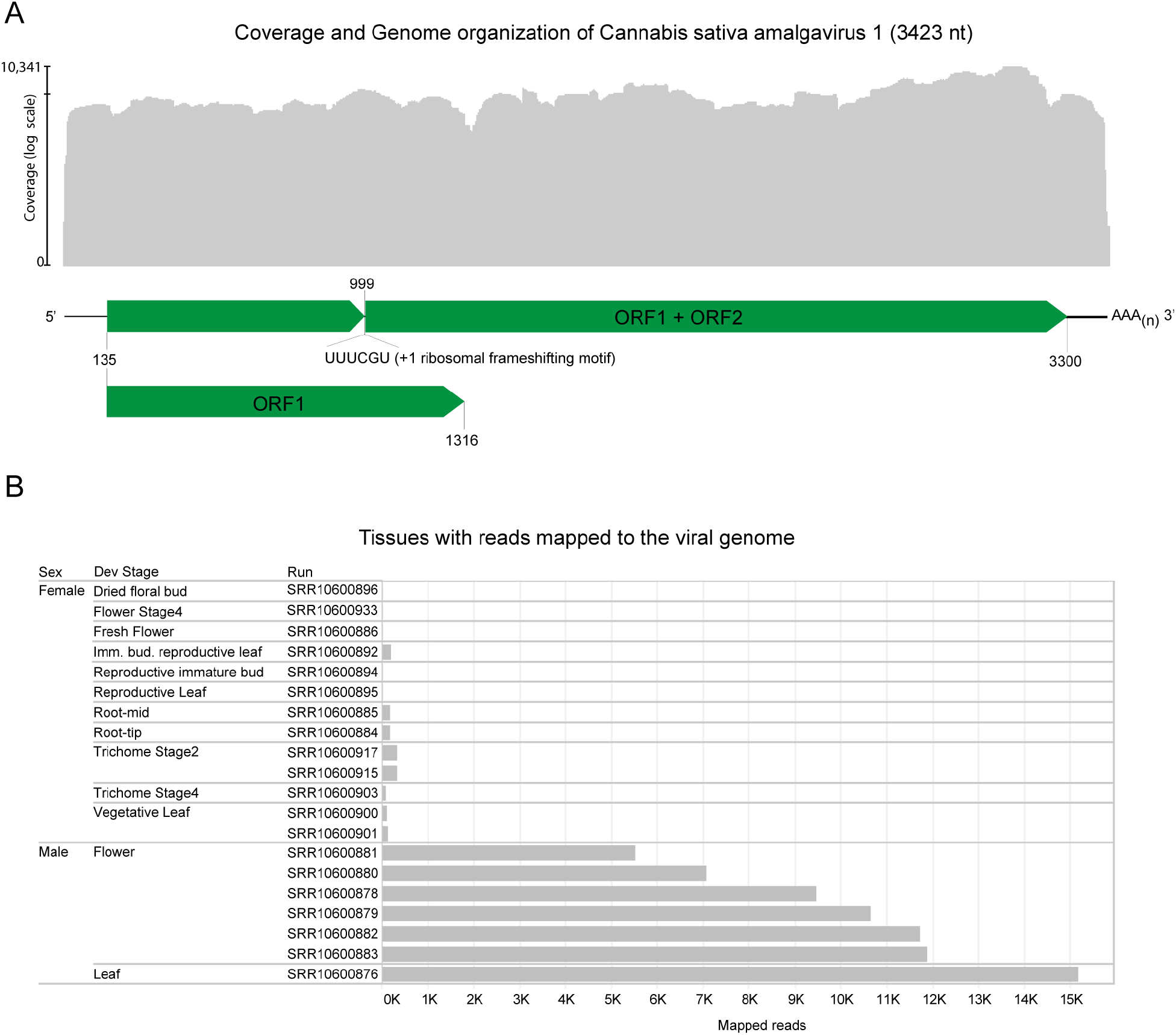
Genome organization, coverage, and detection patterns in distinct tissues of the Cannabis sativa amalgavirus 1 (CSA1) (Genbank accession: MZ707304). **a** Schematic representation of the genomic organization of CSA1 overlaid by its coverage. ORFs are indicated by green arrows whose direction represents the orientation of the transcript (5’-3’). ORF1 encodes a product of unknown function while ORF 1 + 2 encodes a protein function with polymerase domains. The numbers at the ends indicate the position of each ORF’s first and last nucleotide, respectively, and the internal number of the ORF 1 + 2 shows the nucleotide position in which frameshifting occurs, whose motif sequence is amplified. Poly A tail is represented by AAA(n). **b** Tissues specification with the number of reads mapped to the viral genome. In total, it used around 70,000 reads.

Using the criterion of a new species demarcation for the genus Amalgavirus, a divergence height than 25% in the RNA-dependent RNA polymerase (RdRp) amino acid sequence, and differences in the natural hosts’ range (Sabanadzovic and Tzanetakis, 2013), we proceed to the phylogenetic analysis. The RdRp ORF translation of the new Cannabis sativa amalgavirus 1 (CSA1) was compared using the BLASTp tool, 98 related protein sequences were selected, sequences less than 765 amino acids and repeated species were excluded, remaining 43 sequences. We performed a multi-alignment using the MAFFT method (Katoh et al., 2002) v7.450. The phylogenetic relationship was measured using Fasttree (Price et al., 2009) with nodes determined using the Shimodaira–Hasegawa-like approximate likelihood ratio test (SH-aLRT) (Guindon, et al., 2010), and a phylogenetic tree figure was created using FigTree. The pairwise matrix was generated using the virus classification tool SDT (Muhire et al., 2014) (Figure 2). Therefore, the significant divergence in the RdRp ORF and the host qualifies this new sequence as a new amalgavirus species infecting cannabis.

**Fig. 2.**
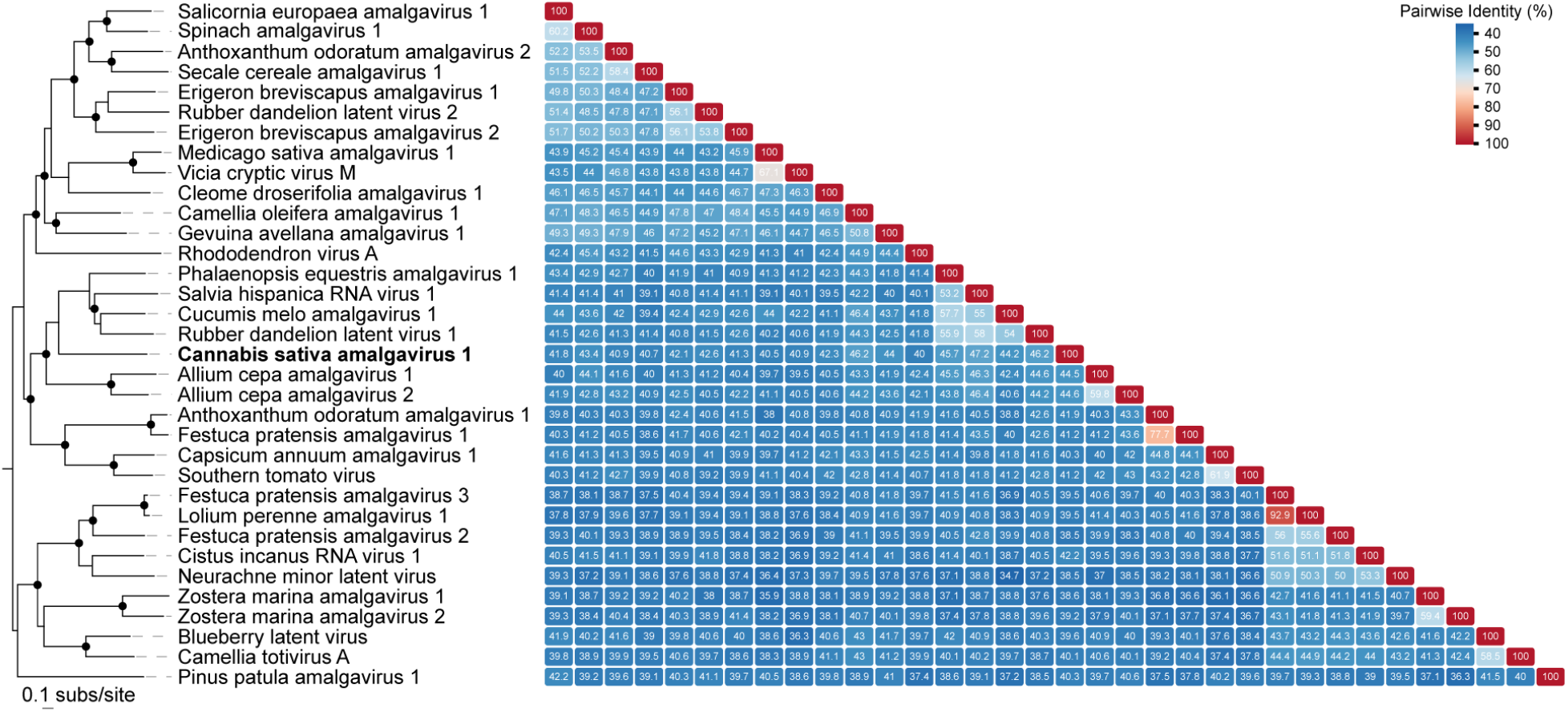
Phylogenetic and similarity analysis based on RNA polymerase amino acid sequences. The phylogenetic tree was created with members from the amalgavirus genus. Sequences were aligned using MAFFT [Katoh et al., 2002], and the maximum likelihood tree was inferred with FastTree [Price et al., 2009]. The black circles represent nodes with a Shimodaira-Hasegawa ≥ 0.9. Cannabis sativa amalgavirus 1 (CAS1) is bolded. The pairwise matrix was generated with the SDT. The color gradient goes from blue to red, indicating the lowest to highest identity values. We adopted the ICTV criteria for the distinction of species proposed for the genus Amalgavirus: Divergence of the polymerase amino acid sequence larger than 25% and differences in the range of natural hosts. All amalgavirus genome sequences available on GenBank until July 15, 2021, were used.

The recent progress in nucleotide sequencing and dataset assembling has generated massive data quantities, much more than our analyzing and annotation capacity. This free and unanalyzed data permits data mining approaches. Indeed in this work, we described the complete genome sequence of a novel amalgavirus in the C. sativa transcriptome. The Cannabis sativa amalgavirus 1 (CSA1) is the first amalgavirus associated with any Cannabis species. Despite the necessity of other bioinformatics resources, investigation and experimental data, its genomic sequence may assist evolution studies and analysis of the possible relationship between new viruses and pathologies that affect *C. sativa*, mainly in the organs where the sequence was detected.

---

**Supplementary Table 1.**

| **Acession Number** | **Tissue-Replicate** | **Total number of reads (Millions)** | **Mapped reads** |
| --- | --- | --- | --- |
| SRR10600883 | Male Flower-Rep1 | 47.1 | 11753 |
| SRR10600882 | Male Flower-Rep2 | 35.1 | 11766 |
| SRR10600881 | Male Flower-Rep3 | 20.3 | 5494 |
| SRR10600880 | Male Flower-Rep4 | 32.2 | 7099 |
| SRR10600879 | Male Flower-Rep5 | 11.8 | 10685 |
| SRR10600878 | Male Flower-Rep6 | 33.2 | 9456 |
| SRR10600876 | Male Leaf | 46.9 | 15269 |

